# Robust RNA Secondary Structure Prediction with a Mixture of Deep Learning and Physics-based Experts

**DOI:** 10.1101/2024.09.18.613732

**Authors:** Xiangyun Qiu

## Abstract

A mixture of experts (MoE) approach is developed to mitigate poor out-of-distribution (OOD) generalization of deep learning (DL) models for single-sequence-based prediction of RNA secondary structure. The main idea is to use DL models for in-distribution (ID) test sequences to take advantage of their superior ID performances, while relying on physics-based models for OOD sequences to ensure robust predictions. One key ingredient of the pipeline, named MoEFold2D, is automated ID/OOD detection via consensus analysis of an ensemble of DL model predictions without accessing training data during inference. Specifically, motivated by the clustered distribution of known RNA structures, a collection of distinct DL models is trained by iteratively leaving one cluster out. Each DL model hence serves as an expert on all but one cluster in the training data. Consequently, for an ID sequence, all but one DL model makes accurate predictions consistent with one another, while an OOD sequence yields highly inconsistent predictions among all DL models. Consensus analysis of DL predictions categorizes test sequences as ID or OOD. ID sequences are then predicted by averaging the DL models in consensus, and OOD sequences are predicted using physics-based models. Instead of remediating generalization gaps with alternative approaches such as transfer learning and sequence alignment, MoEFold2D circumvents unpredictable ID-OOD gaps and combines the strengths of DL and physics-based models to achieve accurate ID and robust OOD predictions.

## 1. INTRODUCTION

RNA, in contrast to DNA and protein, exhibits the unique capacity of dual genetic and non-genetic roles, culminating in the hypothesis of a primordial RNA world (Higgs and Lehman, 2015). Whether in its coding or non-coding roles, RNA executes biological functions through its physical embodiment, a string of nucleotides, governed by universal sequence-structure-interaction principles (Brion and Westhof, 1997; Chen, 2008). Therefore, understanding RNA structure, akin to the protein structure problem, is of central importance for the field of RNA biology. This has led to tremendous interest and progress in elucidating RNA structures with a diverse array of experimental and computational techniques (Cruz, et al., 2012; Das, et al., 2023; Zhao, et al., 2021). As base pairing and stacking dominate the energetics of RNA folding, the pattern and extent of base pairing (i.e., RNA secondary structure) defines the stem-loop motif fundamental to RNA function. This study focuses on single-sequence-based prediction of RNA secondary structure with deep learning (DL) models.

Single-sequence-based (i.e., de novo) DL models aim to learn the sequence-structure mapping of RNA directly from training data. Being highly over-parameterized, DL models are immensely expressive and able to capture the intricate dependencies at very fine-grained levels, as well as pick up various biases and idiosyncrasies present in the training data. These advantages have led to far superior performances of DL models over alternative de novo methods on test RNA samples within the training data distribution (Zhang, et al., 2023). In contrast, non-DL de novo methods typically seek to optimize a score function over all possible base-pairing configurations. The score for a candidate structure can be calculated as its physical free energy with experimentally derived parameters, or as a statistical likelihood with data-derived parameters (Andronescu, et al., 2014) obtained via traditional machine learning (ML) or related techniques (Zhao, et al., 2021). Compared with ML methods, de novo DL algorithms employ orders of magnitude more parameters and are usually trained end to end. As such, DL models have attained state-of-the-art performances and emerged as a very promising solution for de novo prediction of RNA secondary structure (Franke, et al., 2023; Mao, et al., 2022; Wang, et al., 2019; Zhao, et al., 2021).

Instead of operating on a single sequence, another class of predictive models relies on multiple-sequence alignments (MSA) to infer RNA secondary structure from co-evolutionary correlations (Seetin and Mathews, 2012; Zhang, et al., 2023). Often referred to as comparative sequence analysis, hand-crafted algorithms such as mfDCA (Morcos, et al., 2011), PLMC (Hopf, et al., 2017) and RScape (Rivas, et al., 2017) use covariance-based models to predict conserved base pairs. DL models have also been applied to learn from RNA MSAs (Singh, et al., 2021; Sun, et al., 2021; Zhang, et al., 2023). These MSA-based approaches are generally more accurate and robust than single-sequence-based models. For example, recent work showed that MSA-based DL models outperform minimum free energy (MFE) and de novo DL models on new RNA folds (Lang, et al., 2023), though the absolute performances still have much room to improve on new folds defined by the authors, e.g., sequence-averaged F1 scores slightly below 0.6 or 0.5 for models with or without MSA inputs, respectively. Nonetheless, MSA-based structure predictions usually entail high computing cost, because large sequence databases need to be stored and searched, and deep alignments can take hours. The advent of larger databases and more sophisticated pipelines has shown to improve the quality of MSAs but led to even slower MSA generation (Chen, et al., 2024; Zhang, et al., 2022). Conversely, de novo DL methods offer the advantages of rapid prediction speed while maintaining competitive accuracy.

An inherent shortcoming of de novo DL methods, stemming from their sole reliance on training data for learning, is poor out-of-distribution (OOD) generalization. This is manifested as substantial drops in performance for samples that fall outside the training data distribution. Several groups have reported the failure of de novo DL models in cross-family tests, where a specific RNA family (e.g., tRNA) is excluded from the training set but used for testing (Flamm, et al., 2022; Qiu, 2023; Szikszai, et al., 2022). Note that the RNA family here denotes a functional annotation that is different from Rfam’s definition of family based on sequence and structure homology for covariation analysis (Kalvari, et al., 2020). Critically, de novo DL models perform significantly worse than MFE-based models in such cross-family tests. Our further analysis showed that the generalizability of de novo DL models degrades appreciably when the test sample falls below 80% similarity level to the training set (Qiu, 2023). It is worth noting that Lang et al. demonstrated that de novo DL models can achieve comparable performances with MFE models on new RNA folds, with F1 scores around 0.45 (Lang, et al., 2023). We speculate that new RNA folds are less distant than cross-family sequences, as much lower scores are obtained in cross-family tests, e.g., F1 scores ∼ 0.27 for signal recognition particle and 0.36 for transfer-messenger RNA (Qiu, 2023). In short, OOD generalization, particularly at the cross-family level, remains a major challenge for de novo prediction with DL models.

Consequently, various approaches have been proposed to mitigate overfitting and improve OOD generalization of de novo DL models. For example, regularization techniques such as dropouts and weight decays are commonly employed, SPOT-RNA uses model ensembles (Singh, et al., 2019), and MXfold2 further introduces thermodynamic regulation (Sato, et al., 2021). However, while reducing the performance gaps between training and test sets, these strategies often degrade model performance and, importantly, do not fundamentally resolve ID-OOD generalization gaps—they cannot equip de novo DL models with the folding principles of unseen RNA families that are unknown. One effective approach is the aforementioned data augmentation with MSAs (Zhang, et al., 2023), but it is time-consuming and requires high quality MSAs that are not always possible. An alternative way of data augmentation is through representation learning with large language models (LLM). RNA LLMs can be trained in a self-supervised manner on vast amounts of RNA sequences (e.g., from RNAcentral (The, et al., 2017)) to learn semantic, contextual, and distributional information from the entire biological sequence space. Several studies have finetuned RNA LLMs for RNA secondary structure prediction (Chen, et al., 2022; Wang, et al., 2023) with state-of-the-art performances, though it is unclear whether LLM parameters were also trained during fine-tuning. For instance, one recent study reported superior cross-family test results than de novo DL models (Penić, et al., 2024), while revealing inconsistent cross-family generalizations among other existing LLM-based models. Moreover, it remains to be established whether LLM-augmented DL models can eliminate the gaps between training and test performances entirely, which are actual measures of OOD generalization and usually not reported. In essence, while ID-OOD generalization gaps can be mitigated, they are likely to persist when the learning process is (partly) driven by data.

In addition to the inherent limitations of data-driven DL algorithms, the scarcity of known RNA structures is another fundamental cause of the ID-OOD performance gaps. High resolution atomic structures are the best sources for extracting RNA secondary structures, but the number of RNA structures in the protein data bank (PDB) is rather small, fewer than one thousand post redundancy removal (Szikszai, et al., 2024), and the distribution of known structures is largely clustered around a handful of well-studied systems such as tRNA and ribosomal RNA (Schneider, et al., 2023). As a result, existing training data mainly comprise secondary structures inferred through homology-based computational methods. Rfam curates the most comprehensive database of this kind (Kalvari, et al., 2020). Despite being much larger than the quantity of RNA entries in PDB, Rfam collections are highly uneven in terms of length distribution and coverage of RNA functions. The quality of homology-based structures also depends on the quantity and quality of homologous sequences. Therefore, the current state of data quantity, distribution, and quality presents another major obstacle to de novo predictive algorithms solely driven by data.

Here we propose a mixture-of-experts (MoE) approach that integrates excellent performances of de novo DL models for in-distribution (ID) sequences and robust performances of MFE models for OOD sequences. A critical component of our method is automated OOD detection without requiring sequence alignment against the training set. This is realized through consensus analysis of an ensemble of DL models trained via a leave-one-cluster-out (LOCO) approach. Specifically, with the training set grouped into N dissimilar families (or clusters), N DL models are trained by iteratively excluding a single cluster from the training data, resulting in N independent LOCO models, each of which has been exposed to N-1 clusters. During inference, for an ID test sequence, N-1 models are expected to yield highly consistent predictions, while an OOD sequence will result in N dissimilar predictions. Consequently, OOD detection can be automated via consensus analysis of the N predictions. The final output for an ID sequence is the average of the N-1 consistent models, and, as an added benefit, the excluded cluster of the outlier model identifies its cluster (or family) membership. On the other hand, OOD sequences are predicted with an MFE model, LinearFold with thermodynamics parameters (Huang, et al., 2019). In this study, we demonstrate the utility of this workflow, named MoEFold2D, with a medium-sized de novo DL model, SeqFold2D with 960K trainable parameters (Qiu, 2023), and a customized training set comprising nine RNA clusters. Furthermore, we expect this type of MoE approach to be applicable to other scientific domains where only scarce amount of data with clustered distribution are available.

## 2. METHODS

### 2.1 Datasets

Our objective is to create a dataset with a relatively small number (e.g., fewer than a dozen) of distinct clusters, so as to keep the number of LOCO DL models low and maintain a manageable computational cost. We start with two commonly used datasets, ArchiveII (Sloma and Mathews, 2016) and StrAlign (Tan, et al., 2017), collectively containing ten functional types (e.g., tRNA and RNase P). Note that RNA types are used in place of RNA families hereafter to avoid ambiguities with Rfam’s family annotation. We further remove RNA samples with lengths exceeding 600 nucleotides and reduce sequence redundancy levels below 90% identity using CD-HIT-EST (Fu, et al., 2012). The two together result in a total of 8037 samples across nine types. Next, we extract 4837 samples from the bpRNA dataset (Danaee, et al., 2018) that fall within the nine types based on provided metadata, followed by the same length and redundancy processing steps as described above. All samples are then pooled together and a final de-redundancy step with CD-HIT-EST leads to a collection of 9995 samples at the 90% sequence identity level. This collection, referred to as ClustRNA2D, is the main training dataset used in this study. Notably, the populations of the nine RNA types are highly imbalanced, in addition to their widely different length distributions (see SI Figs. 1&2 for details).

Each RNA functional type in the ClustRNA2D dataset is treated as a distinct cluster in this study. Such choice is validated by cluster analyses of pairwise similarities obtained from sequence and structure alignments. Six different RNA alignment programs, LaRA 2 (Winkler, et al., 2022), Foldalign (Sundfeld, et al., 2015), LocARNA (Will, et al., 2012; Will, et al., 2007), RNAforester (Höchsmann, et al., 2003; Lorenz, et al., 2011), Gardenia (Blin, et al., 2010), and RNAdistance (Lorenz, et al., 2011), are used to obtain the pairwise similarity matrices (9995×9995) for the entire ClustRNA2D dataset. The first three alignment programs are provided with RNA sequences only, the next two are provided with both sequences and secondary structures in the dot-bracket notation (dbn) format, and the last (RNAdistance) takes dbn structures as sole inputs. For each aligned RNA pair, we define two similarity scores: (a) seqSim, defined as the percentage of identical residues relative to the mean length of the RNA pair, and (b) dbnSim, defined as the percentage of identical dbn symbols (only for RNAforester, RNAdistance, and Gardenia). For similarity-based clustering, the OPTICS algorithm is chosen due to its low sensitivity to varying sample densities. Outliers are assigned into their nearest clusters based on averaged distances to clusters.

### 2.2 DL Model Architecture and LOCO Training

The SeqFold2D DL model (Qiu, 2023), developed by us for studying DL generalization, is chosen for this study. Its architecture and training protocol follow our previous work, as provided in SI Section 2. Briefly, SeqFold2D is a de novo predictive model taking single RNA sequences as the only inputs without data augmentations of MSA or LLM representations. SeqFold2D is also free of post-processing steps such as enforcement of canonical pairs, sparsity, or thermodynamic consistency. The main trunk of SeqFold2D consists of a sequence module (transformer and long-short-term memory, LSTM) and a pair module (2D convolution), and the model output is the 2D base pairing probability matrix (PPM). For this study, a medium-size SeqFold2D model with 960K trainable parameters is selected. This model was shown to achieve competitive performances against existing DL models (Qiu, 2023). SeqFold2D is thus representative of single-sequence-based DL models, and we choose not to include detailed performance comparisons against other DL models in this study.

In the MoEFold2D workflow, each DL model is trained on all but one cluster in the training data. With N=9 RNA clusters in the ClustRNA2D dataset, nine SeqFold2D models are trained independently by iteratively leaving one cluster out of the training set for each model. This results in nine DL models, referred to as the LOCO ensemble.

### 2.3 MoEFold2D Prediction Pipeline

When predicting RNA secondary structures, the LOCO ensemble is expected to exhibit distinct behaviors for ID and OOD test samples. For a test sample belonging to one of the N clusters in the training data, we expect poor performance from only one of the N LOCO models—the one trained with the specific cluster left out—while the other N-1 predictions are expected to be accurate and highly similar to each other and the ground truth. This scenario is readily discernable through cluster analysis of the N LOCO predictions, signified by a single cluster with N-1 similar predictions and one outlier. The N-1 predictions in consensus are then averaged to produce the final prediction for the ID sequence. An added benefit is the prediction of the cluster membership (or RNA type) of the ID sequence, which corresponds to the cluster excluded during training of the outlier model.

Conversely, if the test sample does not belong to any of the N clusters in the training data, we expect all N models to yield poor and dissimilar predictions. Clustering of the LOCO predictions would fail to identify a single cluster of N-1 similar predictions and one outlier. In this scenario, none of the DL models can be trusted, and an MFE-based model is adopted as the predictor. Here we opt for LinearFold (Huang, et al., 2019) for its linear dependence of computing time on the RNA sequence length and competitive performances against other MFE-based methods such as RNAstructure (Reuter and Mathews, 2010) and RNAfold (Lorenz, et al., 2011).

### 2.4 Evaluation Metrics

We choose F1 score as the primary metric for assessing model performance due to its ability to effectively balance precision and recall in classification tasks (Mathews, 2019). In binary classification, F1 score is defined as 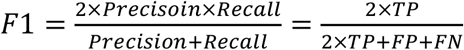 where Precision is TP/(TP+FP), Recall TP/(TP+FN), TP the number of true positives, FP false positives, FN false negatives, and TN true negatives. The TP, FP, and FN values are computed by comparing the predicted pairing probability matrix, after discretization with a threshold of 0.5, with the ground truth matrix of RNA secondary structure. When averaging F1 scores across a dataset, we employ a sequence-wise approach rather than averaging at the cluster or family level.

## 3. RESULTS

### 3.1 Cluster Analysis of the ClustRNA2D Dataset

RNA functional types represent categories of RNA molecules characterized by high levels of sequence and structure conservation, reflecting the fundamental coupling between sequence, structure, and function. Each RNA type thus exhibits distinct sequences and structures, making cross-type validation a common practice for benchmarking OOD performances of predictive algorithms. In this study, we adopt RNA types as unique clusters and further verify their congruence with similarity-based clusters. Our analysis reveals that, for both seqSim and dbnSim scores, pairwise similarity matrices show noticeably higher intra-type similarities compared to inter-type values, as expected (SI Fig. 3). Utilizing the OPTICS algorithm for cluster analysis, we find that the dbnSim scores derived from RNAforester yield the most consistent clusters aligned with the known RNA types. As depicted in Fig. 1, each similarity-based cluster predominantly comprises RNAs from a single type, with minimal “spillover” from other types. Notably, two RNA types, 16S rRNA and SRP, are split into two clusters each, possibly due to substantial intra-type heterogeneity suggested by their non-Gaussian-like length distributions (see SI Fig. 2). Additionally, the two least populous RNA types, 23S rRNA (20 samples) and TERC (30 samples), fail to form distinct clusters and are identified as outliers, which are assigned to closest existing clusters. It is possible to cluster them successfully using OPTICS parameters allowing for low-density clusters, but such clustering also results in frequent splits of other RNA types and numerous clusters with small sizes (see SI Figs. 3&4 and associated text for details).

**Figure 1.**
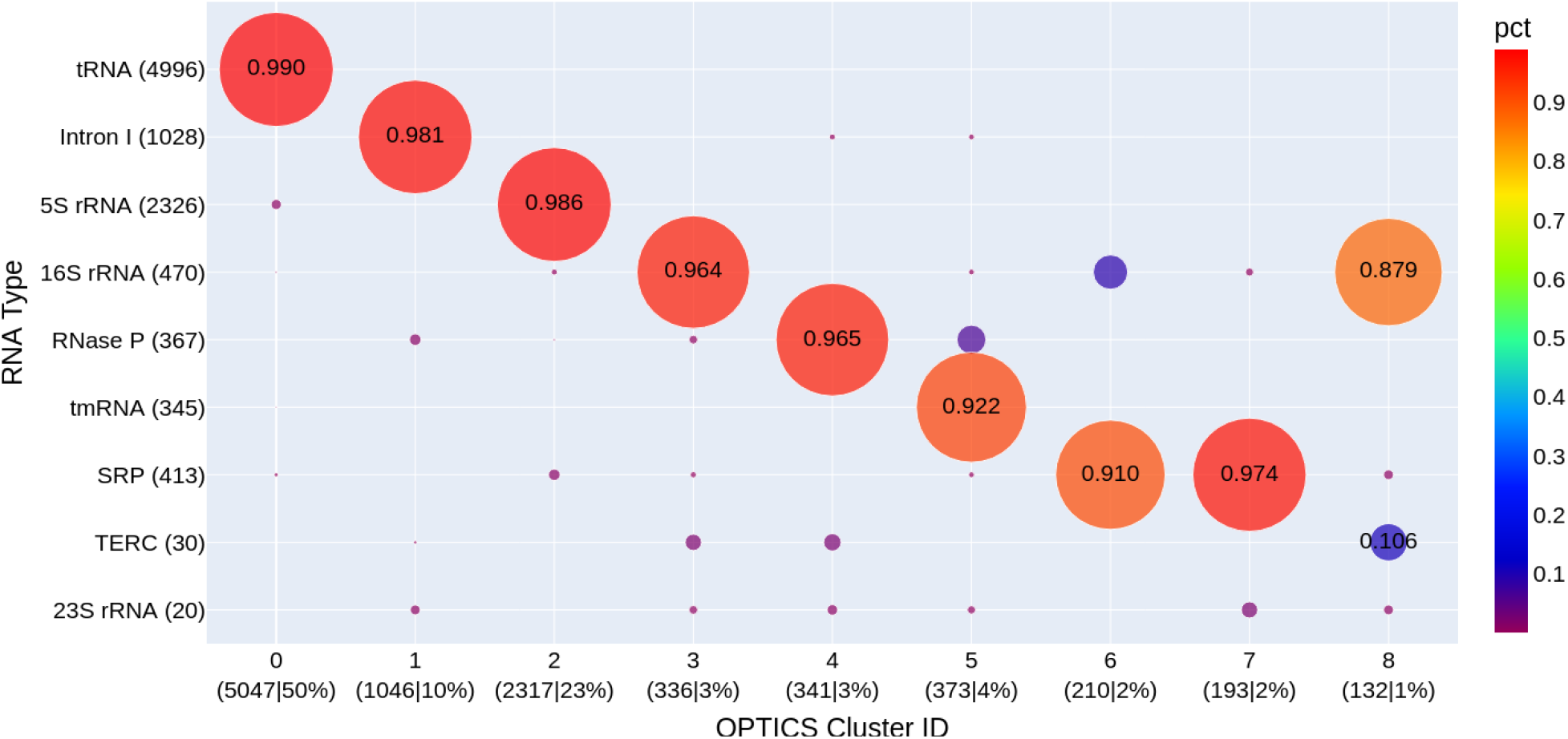
The composition of RNA functional types within each similarity-based cluster identified by the OPTICS algorithm. The size of each circle scales with the count of samples normalized by the size of its parent cluster, with its fractional value shown at the center of the circle. The total number of samples in each RNA type or cluster is given in the tick label within parentheses. The abbreviations of RNA types are, rRNA: ribosomal RNA, tRNA: transfer RNA, Intron I: group I intron, tmRNA: transfer messenger RNA, SRP: signal recognition particle, and TERC: telomerase RNA component. It is important to note that these RNA types, rather than OPTICS clusters, are used as individual clusters for training deep learning models.

Taking these factors into consideration, our observations assert that each RNA type comprises a group of closely related RNA molecules distinct from all other types. However, large differences in densities and non-uniform distributions among RNA types make it rather difficult to achieve perfect cluster vs. type congruence with a single set of algorithmic parameters. In sum, our cluster analyses validate RNA functional types as standalone clusters, while acknowledging substantial variations in cluster sizes and intra-cluster distributions.

### 3.2 Performances of Individual LOCO Models

Here we assess how the performances of invididual LOCO models are influnced by the exclusion of a specific RNA type from the seen set, as well as varied data redundancy levels and type-based upsampling of the seen set. N.B. clusters and types are used exchangeably. ClustRNA2D is first split into one seen set and one unseen set. The unseen set also serves as an ID test set, referred to as TS80, obtained by reducing the sequence redundancy level of ClustRNA2D to 80% (via CD-HIT-EST) and drawing out 15% through cluster-stratified, random sampling.

Then, a seen set for training and validation is obtained by excluding samples in ClustRNA2D that have sequence identity above 80% against TS80. The seen set, referred to as NR90, thus has an intra-set sequence redundancy level similar to ClustRNA2D (i.e., 90%). The rationale for a seen set with higher redundancy than a typical 80% is to expose DL models to more diverse samples within the same overall distribution. However, this is at the expense of even more imbalanced size distribution among the clusters, as the more populous clusters typically have higher intracluster densities in the original dataset. In order to investigate the effects of varied cluster sizes and redundancy levels, we consider several additional configurations of the NR90 set: a) NR90-UP, by up-sampling all clusters to match the size of the largest cluster; b) NR80, by further reducing NR90 to 80% redundancy with CD-HIT-EST; and c) NR80-UP, by up-sampling the NR80 set to have the same size for all clusters. For each of these seen set configurations, we train a total of ten SeqFold2D 960K models, nine LOCO models and one with the entire seen set. Each seen set is further randomly split into a training set for updating trainable parameters and a validation set for early stopping, and TS80 is the common test set.

The four seen sets (NR90, NR90-UP, NR80, and NR80-UP) yield distinctive model performances, as depicted in Fig. 2A. Notably, up-sampling results in nearly identical *training set F1 scores* across all RNA clusters, irrespective of their original cluster sizes. This stands in stark contrast to the degradation of training performances observed for under-represented clusters without up-sampling. However, consistent training performances on up-sampled clusters do not transfer to the validation or test sets, where larger train-validation gaps are evident for clusters with smaller sizes, and the train-test gaps generally widen even further. While up-sampling levels up the training performances across all clusters, it fails to achieve similar improvements on test samples, limiting its effectiveness in mitigating cluster size imbalances.

**Figure 2.**
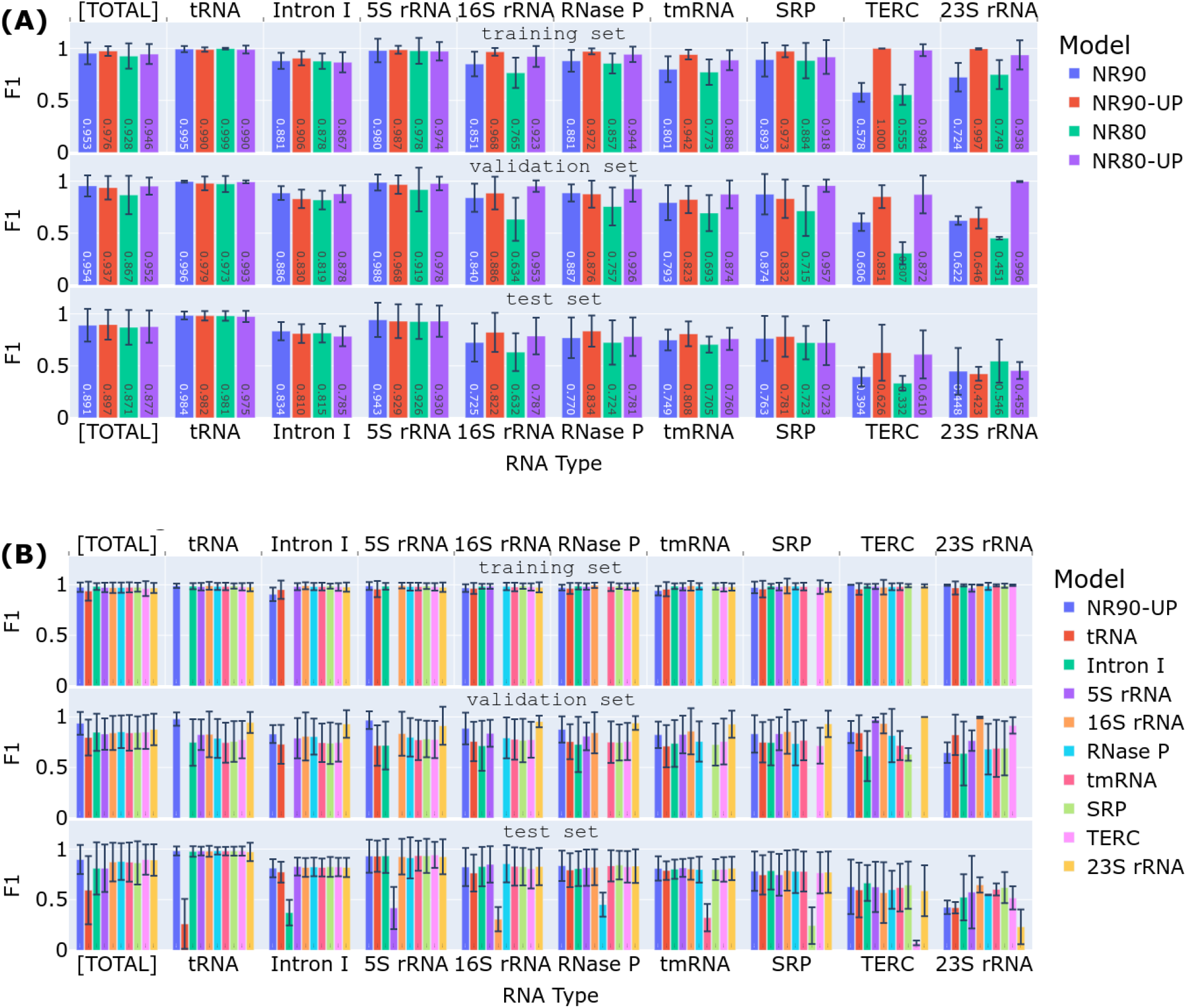
The performances of SeqFold2D models trained on different data configurations. In each panel, the averaged F1 scores are displayed for the respective training (top), validation (middle), and test (bottom) sets. The training and validation sets are random splits of the seen set, which is detailed below. The test set, TS80, is common for all models. For each set, the scores are presented for the entire set, denoted as [TOTAL], and for its constitutive RNA type labeled along the x axis. (A) SeqFold2D models trained on four seen sets: NR90, NR90-UP, NR80, and NR80-UP, respectively. Each model is identified by its seen set, as specified in the legend. (B) SeqFold2D models trained via the LOCO strategy on the NR90-UP seen set, with the left-out cluster/type indicated in the legend. A SeqFold2D model trained with no cluster left out is shown as a reference, denoted as NR90-UP. Note that each LOCO model (all except NR90-UP) excludes its respective left-out cluster from both its training and validation sets, resulting in empty bars in corresponding positions.

Upon closer examination of the common test set, TS80, we observe slightly higher F1 scores for the NR90 and NR90-UP sets compared to NR80 and NR80-UP for the entire TS80 set (denoted as [TOTAL] in Fig. 2). For individual RNA clusters, the under-represented clusters (e.g., TERC and 16S rRNA) benefit more from higher redundancy (e.g., NR90 vs. NR80) or up-sampling (e.g., NR90-UP vs. NR90) of the training set. However, it is important to note that the improvements are small, and under-represented clusters still exhibit rather low F1 scores, particularly for the case of sparsely populated 23S rRNA with very poor outcomes from all four seen sets. Overall, our observations indicate that training with a seen set featuring higher redundancy or up-sampling can enhance model performances, albeit to a modest degree and dependent on the specific data distributions. Consequently, NR90-UP is used as the seen set for the results presented hereafter, while noting that the uses of the other sets yield comparable outcomes.

The presence of nine RNA clusters/types in the ClustRNA2D dataset enables the training of nine SeqFold2D models using the LOCO strategy. Fig. 2B illustrates their performances when trained with the NR90-UP set, along with a model trained with the entire NR90-UP set for comparison. It is evident again that up-sampling levels up training performances for all clusters, irrespective of their sizes, but remains susceptible to overfitting and poor generalization. Crucially, for every RNA type in the test set (TS80), we observe highly similar F1 scores from all models except one, and the outlier model is the one trained with that specific RNA type left out. Moreover, the outlier model exhibits notably poorer performance compared to the other LOCO models, as expected for ID samples, which is the case for TS80 with respect to NR90-UP. Similar trends are observed on LOCO model ensembles trained on the NR90, NR80, and NR80-UP sets (SI Figs. 7-9). These observations lend strong support for the proposed method of automated ID/OOD detection, where ID samples can be identified by the pattern of a single cluster of N-1 highly similar predictions and one outlier among the N LOCO predictions.

### 3.3 Ensemble-based Prediction of ID Samples with Known RNA Types

With nine LOCO models as shown in Fig. 2B, an ID sample results in eight consistent predictions and one outlier, while an OOD sample yields nine mutually dissimilar predictions. Automated ID/OOD detection can thus be achieved via consensus analysis of LOCO predictions. Beyond ID/OOD classification, the left-out RNA type of the outlier model further predicts the RNA type of an ID sample, whereas “OOD” is assigned to OOD samples. Thus predicted RNA types for the TS80 test set are shown against the ground-truth RNA types in Fig. 3A with NR90-UP as the seen set. Out of the 606 samples in TS80, 576 (95%) are correctly predicted as their ground-truth types, 4 (0.7%) are mis-identified as other types, and 26 samples (4.3%) are labeled as OOD. The overall performance is excellent, and many RNA types are predicted nearly perfectly. Nonetheless, the success rate of ID/OOD detection depends on the RNA type and appears to strongly correlate with the test performance of the specific type. For example, the lowest success rates are observed for 23S rRNA (0%) and TERC (33%) with test F1 scores are far lower than the other RNA types, below 0.6 vs. over 0.9 (Fig. 2B). This correlation is rather healthy as we prefer not to identify a sample with a low F1 score as ID and predict its type (further discussed below). On the whole, the LOCO ensemble proves to be capable of identifying the specific cluster membership for ID test samples.

**Figure 3.**
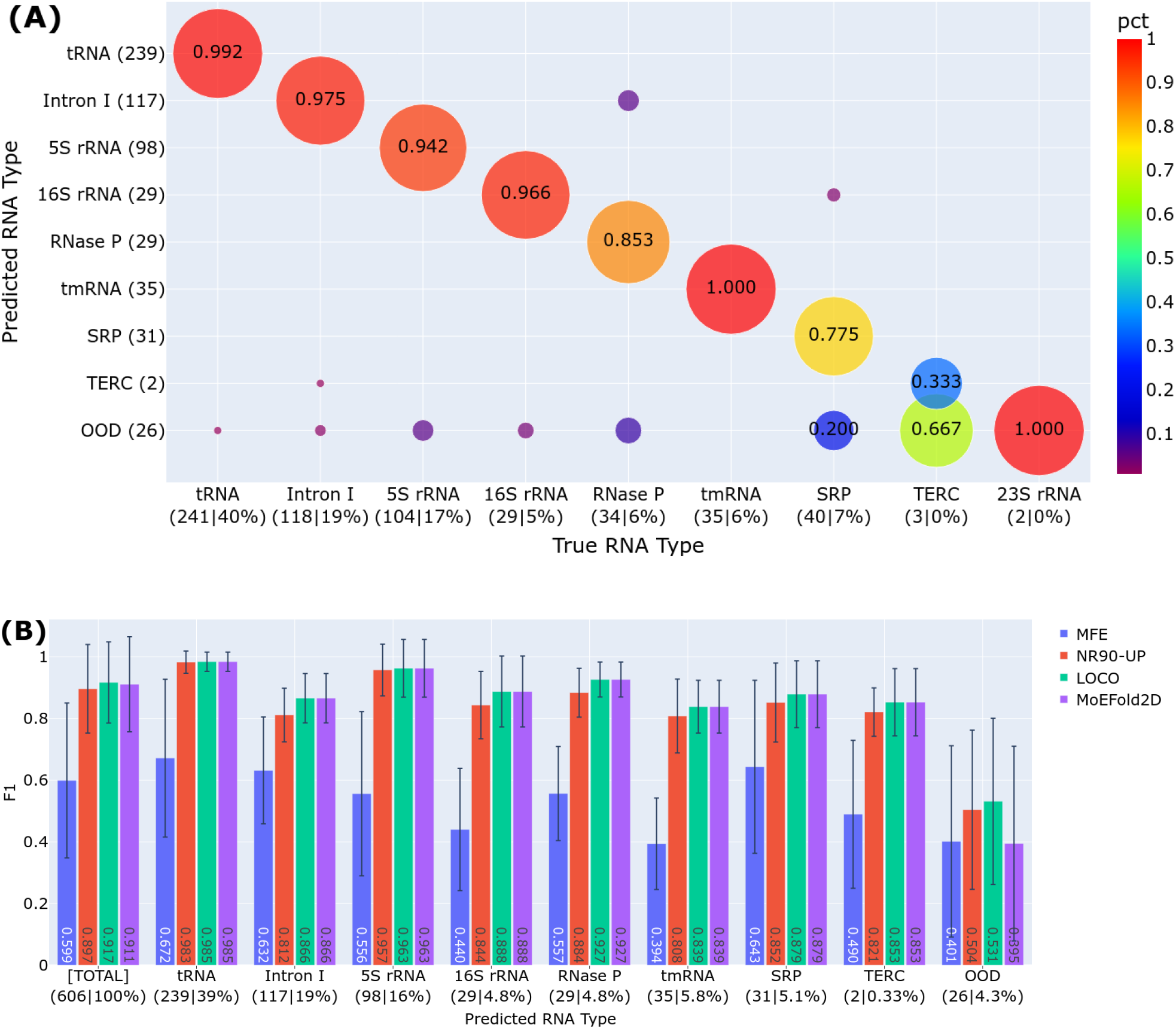
Model predictions on the TS80 test set with known RNA types. (A) Congruence between the predicted (y axis) and ground-truth (x axis) RNA types of the TS80 set. (B) Performance comparisons of different models: MFE, SeqFold2D (NR90-UP), the LOCO ensemble (LOCO), and MoEFold2D. F1 scores are shown for the entire TS80 set ([Total]) and the predicted RNA types.

Importantly, it appears insufficient for ID samples to simply belong to one of the training types to achieve successful RNA type prediction. Instead, the true prerequisite for ID samples is consistent predictions from all but one LOCO models. High consistency among multiple independent LOCO models gives a very high chance for all predictions to be close to the ground truth and hence highly accurate. As such, samples that categorically belong to the training types can still be classified as OOD if the LOCO ensemble makes inconsistent predictions, which are very in fact very likely to be inaccurate. Notably, this consistency-based classification is not a disadvantage but an advantage of the LOCO strategy, as our overarching goal is to use DL models where DL predictions are accurate. Here the consensus analysis presents a practical way to gauge the prediction robustness and accuracy of DL models, going beyond the use of nominal RNA types that are imperfect surrogates for ID/OOD detection.

In the next step of the MoEFold2D workflow, the secondary structures of ID and OOD samples are generated in different ways: ID samples take the average of their respective sets of consistent LOCO predictions while OOD samples are predicted with an MFE-based model (LinearFold). Fig. 3B shows the MoEFold2D performances on the TS80 set, together with the MFE-based predictions of all TS80 set (denoted as “MFE”), the predictions by a single SeqFold2D model trained with the entire NR90-UP set (“NR90-UP”), and the averages of the LOCO ensemble (“LOCO”) where consistent models are averaged for ID samples and all models are averaged for OOD samples as the best guess. As expected from TS80 being an ID test set, DL models (SeqFold2D and LOCO) surpass the MFE model by substantial margins overall and individually for each identified RNA type. It is reassuring that the test samples with successful identification of RNA types all have excellent performances, and the 4.3% samples detected as OOD show much poorer performances in comparison, corroborating the ability of consensus analysis in estimating model accuracies. It is worth noting that the LOCO ensemble further shows a slight edge over SeqFold2D, presumably due to the averaging of multiple LOCO models. In sum, the case study of TS80 as an ID test set lends strong support to the utilities of the MoEFold2D workflow in automated ID/OOD detection, identification of RNA types, and superior performances of ID samples.

### 3.4 Ensemble-based Prediction of OOD Samples with Unknown RNA Types

We next examine the LOCOFold performances on an OOD test set with unknown RNA types. The TS0 set, first compiled by the SPOT-RNA team, is used for this purpose and referred to as bpRNA-TS0. As bpRNA-TS0 is derived from bpRNA with partial overlaps with ClustRNA2D, we further filter out the samples above 80% sequence identity with respect to the entire ClustRNA2D dataset using CD-HIT-EST, obtaining a total of 1118 samples out of 1305 in the original set. Fig. 4 shows the F1 scores on the filtered bpRNA-TS0 set as a whole ([Total]) and grouped by the predicted RNA types. Over 86% of the bpRNA-TS0 set is predicted as OOD, consistent with bpRNA-TS0 being much more distant to the seen set (NR90-UP) than the TS80 set. Evidently, the MFE model substantially outperforms the DL models (NR90-UP and LOCO in Fig. 4) on the OOD samples, as expected from the poor OOD generalization of DL models. Interestingly, on the 14% of bpRNA-TS0 identified as ID, the DL models achieve slightly better performances than the MFE model. By choosing DL for ID samples and MFE for OOD samples, MoEFold2D combines the best performances on both ID and OOD populations, testifying to the robustness of the MoEFold2D workflow on a distant test set.

**Figure 4.**
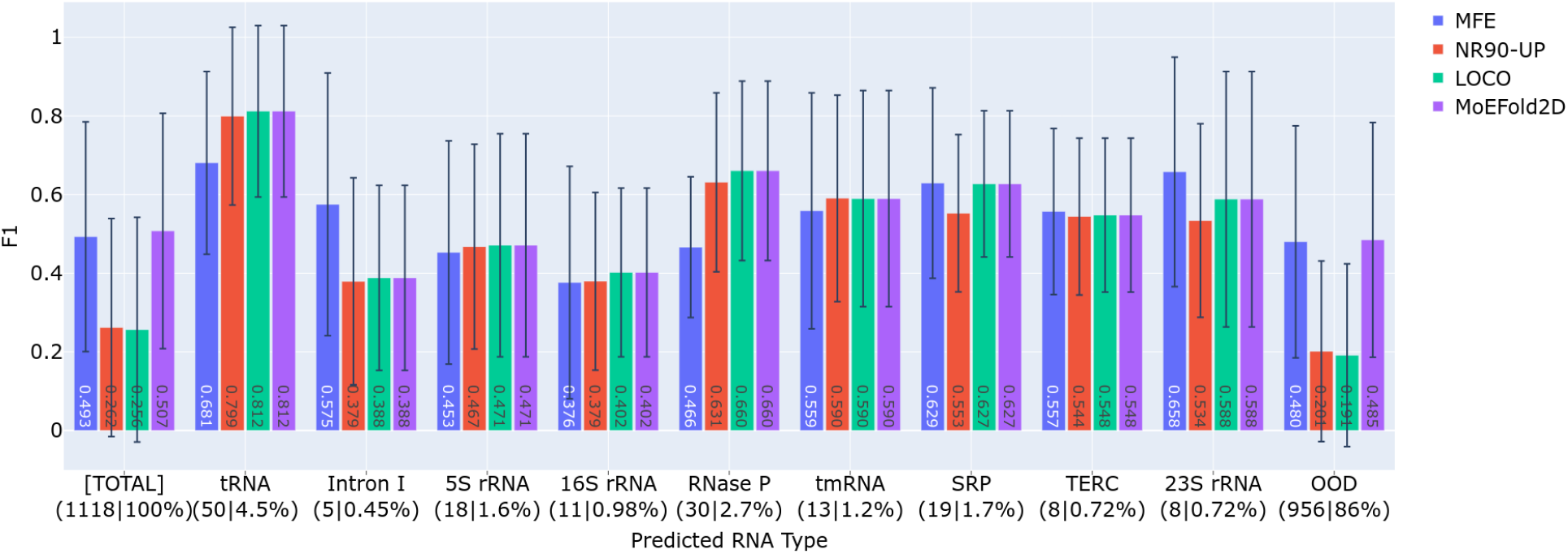
Model performances on the filtered bpRNA-TS0 test set with unknown RNA types. The same four model (groups) as in Fig. 3 are shown here as denoted by the legend. Note that the MFE and MoEFold2D predictions are the same for OOD samples.

The case study of bpRNA-TS0 as a distant test set indicates that the LOCO DL models are only used for predicting a rather small fraction of test samples and mainly serve as the method of automated ID/OOD detection. Despite the reduced utility of DL models, it can be argued that this is the preferred action because DL models cannot be trusted on OOD samples. This case study further demonstrates that the consensus analysis of LOCO ensemble can pick out the samples among a distant set that are likely predicted with high accuracy by DL models, evidenced by the large performance gaps between ID and OOD samples (NR90-UP and LOCO in Fig. 4). This corroborates the advantage of consensus analysis in gauging the prediction accuracy of DL models.

Notably, the absolute performance of the DL models on the ID samples in bpRNA-TS0 is significantly lower than that on the ID samples in TS80, as low as 0.4 for some predicted RNA types. As the consensus analysis clusters together predictions within a radius of 0.2 in F1-score distance from the center, the pair distances within a cluster can range from zero to nearly 0.4. The cluster radius of 0.2 can be tuned up or down to loosen or tighten the consistency threshold, which would also lead to more or fewer samples identified as ID, respectively. A more stringent threshold generally improves the prediction accuracy but can render all samples identified as OOD. Here 0.2 is chosen such that the DL models would still outperform or at least match the MFE model on the ID samples in bpRNA-TS0. Moreover, the pairwise F1 scores used by the consensus analysis contain additional statisticas such as the mean and spread of the scores. We have examined the informative power of such statistics on prediction accuracy but found no satisfactory correlation near the boundary of ID/OOD classification. More work is required to extend the consensus analysis beyond binary ID/OOD classification.

## 4. CONCLUSION

In this study, we present a practical and effective approach to take advantage of superior ID performances of DL models via a LOCO strategy to first realize automated ID/OOD detection. The LOCO method is enabled by the clustered distributions of RNA sequences with reliable knowledge of secondary structures, allowing training of an ensemble of DL models each of which is exposed to all but one data cluster. Consensus analysis of the LOCO ensemble serves to identify ID samples in the test set and further designate their cluster membership without accessing the training data (i.e., in lieu of comparative sequence analysis). With ID samples predicted by averaging the LOCO ensemble, and OOD samples predicted by an MFE model, the MoEFold2D pipeline achieves highly accurate ID and robust OOD predictions. Two case studies with one ID and one OOD test sets demonstrate the efficacy of MoEFold2D in ID/OOD detection and prediction performance.

MoEFold2D can be substantially improved in multiple aspects beyond the proof-of-principles studies presented here. First, expansion of existing clusters and addition of new clusters will improve the performance and scope of the LOCO ensemble. Several clusters in ClustRNA2D are either too small (e.g., 23S rRNA and TERC) or very heterogenous (e.g., SRP and 16S rRNA), and increasing the population and coverage of these existing clusters will boost both prediction accuracy and ID detection for these specific RNA types. Addition of new clusters will broaden the span of ID distributions as well as the utilization of ID predictions of DL models. One possibility is to incorporate the entire Rfam database into ClustRNA2D. However, our preliminary analysis observes the appearance of many small-sized clusters and the merge of some existing clusters (e.g., tRNA and 5S rRNA). Detailed analyses, including data pruning and alternative clustering algorithms (e.g., bpRNA-align (Lasher and Hendrix, 2023) or graph-based GraphClust (Heyne, et al., 2012)), are likely necessary for cluster-based curation of a diverse and large dataset such as Rfam. On the model side, a larger SeqFold2D model may be needed for training a larger number of clusters with increased diversity in order to obtain superior ID performances. The MFE model for OOD samples, currently LinearFold only, can be extended to an ensemble of physics-based models and to make use of the recent update of thermodynamic parameters (Mittal, et al., 2024). Moreover, the consensus analysis can be replaced by a machine learning model or multilayer perceptron trained for ensemble learning. Lastly, the set of LOCO DL models all have distinctive learning dynamics resulting from different left-out clusters, and more fine-grained analysis of the LOCO ensemble predictions (i.e., beyond the consistency analysis based on mutual F1 scores) may shed lights on the uncertainty quantification or structural diversity. Conversely, the premise behind MoEFold2D, poor OOD generalization of DL models, will become obsolete when the body of RNA sequences with known structures eventually covers the explorable RNA space in biology.

Meanwhile, the MoEFold2D pipeline, particularly the LOCO ensemble method, can be readily transferred to other scientific domains with sparse and clustered data distributions, so as to take advantage of the expressive capacity of DL for ID predictions. More broadly, the MOE approach is witnessing a renaissance in the realm of large language and multimodal models where an ensemble of smaller, task-specialized DL models can be trained and run at lower costs and yield competitive performances collectively (Fedus, et al., 2022; Lin, et al., 2024). The MoEFold2D approach herein develops data-specialized DL models, and consensus analysis is used in place of a routing network for ID/OOD detection. Its small mode size and compute requirement (e.g., a single commodity GPU with 12 GB RAM for the SeqFold2D 960K model) further make DL accessible to the broader scientific community for rapid exploration.

## ACKNOWLEDGEMENT

We thank the GW High Performance Computing Facility for free access of CPU and GPU nodes.

## AVAILABILITY

MoEFold2D is open-source software available in the GitHub repository (https://github.com/qiuresearch/MoEFold2D).

