## Supplementary material for "Robust RNA Secondary Structure Prediction with a Mixture of Deep Learning and Physics-based Experts": SI text and figure

Xiangyun Qiu*
Department of Physics, George Washington University, Washington DC 20052

### 1. ClustRNA2D Dataset

**Overview.** The main objective is to enable cluster-based method developments such as the training of deep learning (DL) models via the Leave-One-Cluster-Out (LOCO) strategy described in the main text. A new dataset, named ClustRNA2D, is thus curated by collecting RNA entries with known functional classifications from three databases, ArchiveII (Sloma and Mathews, 2016), StrAlign (Tan, et al., 2017), and bpRNA (Danaee, et al., 2018). The ClustRNA2D functional types (e.g., tRNA, rRNA, and self-splicing introns) are found to be valid cluster labels via clustering analysis based on pairwise sequence and structure similarities. It is worth noting that the RNA types or clusters here are much larger than the RNA families denoted by Rfam which are based on rather strict sequence and structure homology required for covariance analysis. In consideration of the large computational cost of pairwise similarity computation, we exclude RNA sequences with lengths over 600 bases and filter out sequences with redundancy levels above 90% (by CD-HIT-EST (Fu, et al., 2012) with default parameters). Note that each cluster is usually further reduced in redundancy for DL training. Specifically, there are a total of nine RNA functional types whose population and length distributions are shown in Figs. 1&2.

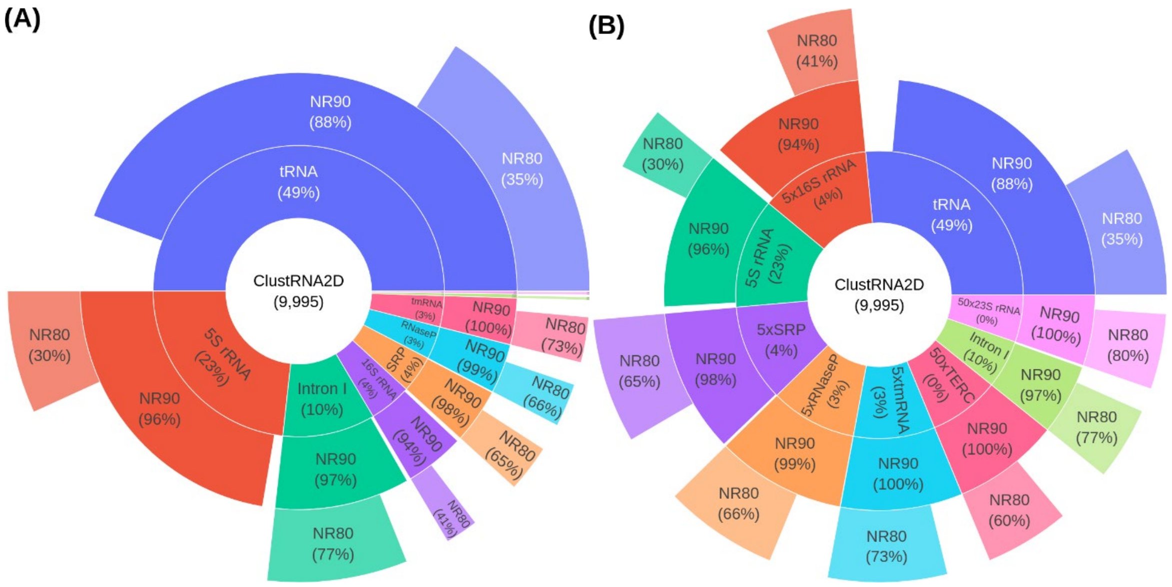


Figure 1 Population distributions of different RNA functional types of the ClustRNA2D dataset at different redundancy levels. For both panels, the first ring shows each RNA type along with its percentage in the entire dataset; NR90 and NR80 denote non-redundancy levels at 90% and 80%, respectively, along with its ratio relative to the original size for each type. Panel (A) shows pies in proportion to the populations of RNA types, while panel (B) scales them for visibility. The abbreviations are, rRNA: ribosomal RNA, tRNA: transfer RNA, Intron I: group I intron, tmRNA: transfer messenger RNA, SRP: signal recognition particle, and TERC: telomerase RNA component.

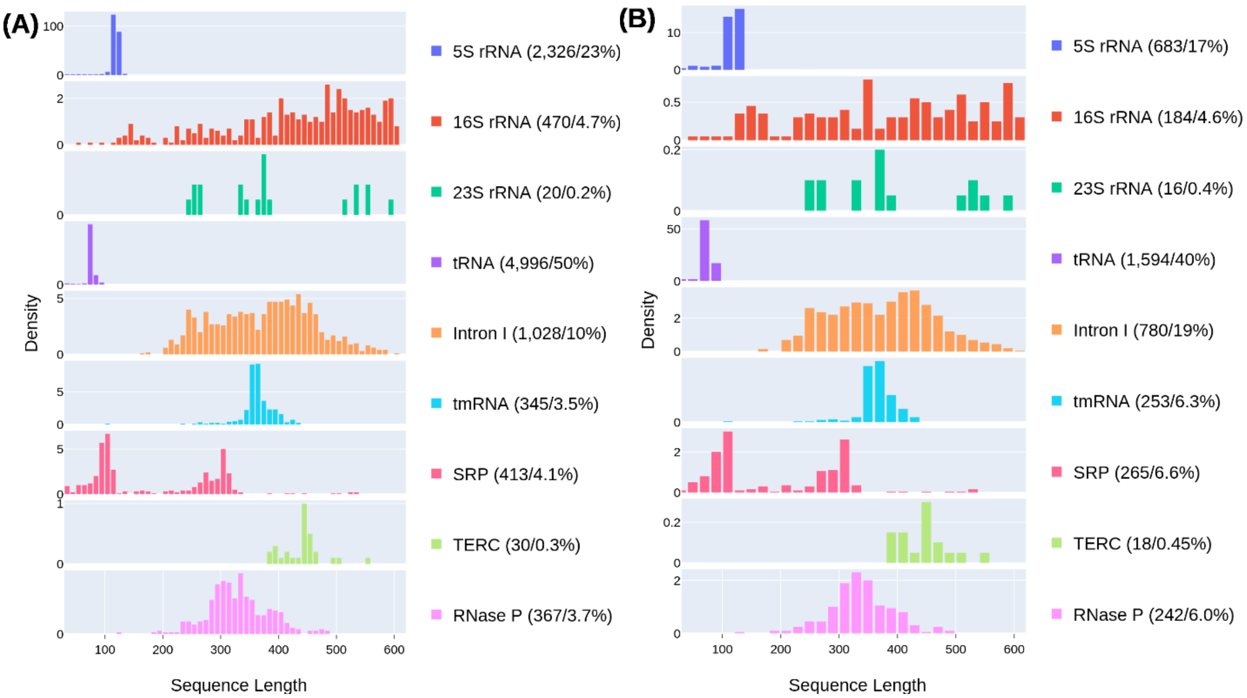


Figure 2 Length distributions of the different RNA functional types in the ClustRNA2D dataset (A) and its subset at the 80% sequence identity level (B). The total population count, and percentage of each RNA type are shown in the legend between a pair of parentheses.

**Cluster analysis.** As our goal is to identify clusters delineated by sequence and structure similarities, we further obtain the pairwise similarity matrix and cluster the sequences with the OPTICS algorithm. Six sequence and structure alignments are tested: LaRA 2 (Winkler, et al., 2022), Foldalign (Sundfeld, et al., 2015), LocARNA (Raden, et al., 2018), RNAforester (Lorenz, et al., 2011), Gardenia (Blin, et al., 2010), and RNAdistance (Lorenz, et al., 2011). Only RNA sequences are provided for the first three programs, though all of them fold and align the pair of sequences internally; the next two programs take both sequences and their ground-truth secondary structures as inputs; the last (RNAdistance) aligns secondary structures only. For each aligned RNA pair, we then compute two types of similarity scores: (a) seqSim, defined as the percentage of identical residues relative to the mean length of the RNA pair, and (b) dbnSim, defined as the percentage of identical dbn symbols (only for RNAforester, RNAdistance, and Gardenia). Examples of the seqSim and dbnSim matrices (9995×9995 for the full ClustRNA2D dataset) are shown in Fig. 3 by grouping RNAs by their functional types. As expected, the similarity scores within each RNA type (along the diagonal) are substantially higher than across RNA types (off diagonal), and the dbnSim scores are generally higher than seqSim as structure is more conserved than sequence. Close inspection further reveals wide-varying intra-type and inter-type similarities that depend on numerous factors such as the size and density of each RNA type and the evolutionary distances between types. For example, tRNA and 5S rRNA are the most similar inter-type pairs and several types (RNaseP, 16S rRNA, and group I intron) display above-average similarities to other types.

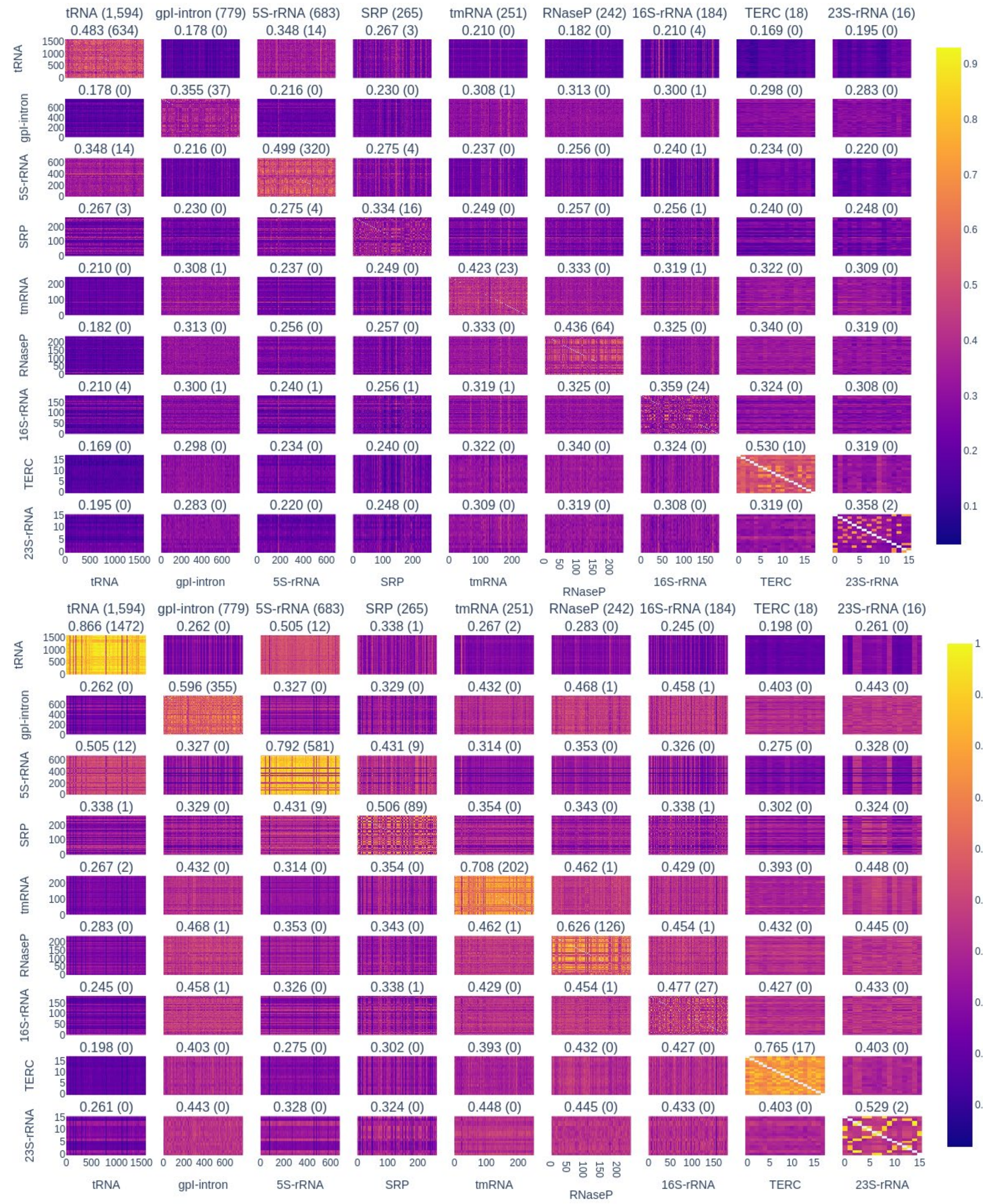


**Figure 3** Heatmaps of the seqSim (top) and dbnSim (bottom) matrices of the ClustRNA2D NR80 dataset. The total count of each RNA type is shown at the top row within the parentheses. The title of each grid shows the average similarity score and the number of unique pairs with values above 0.5 (seqSim) or 0.6 (dbnSim).

Pairwise matrices of the seqSim and dbnSim scores are then clustered with the OPTICS algorithm from the Scikit-learn.cluster module in Python. Outliers found by the algorithm are assigned to the clusters with nearest average distances. Generally, we find significant inconsistency between the seqSim-based clusters and RNA types. Much better congruences are observed between dbnSim-based clusters and RNA types, with RNAforester yielding the best agreement, followed by RNAdistance. It is important to note that clustering outcomes strongly depend on algorithmic parameters, especially minimum similarity and minimum number of samples. Our choices of parameters are guided by the inspection of pairwise similarity matrices (Fig. 3) and prior knowledge of RNA functional types. Complementary to Fig. 1 in the main text, Fig. 4 herein shows how each RNA type is distributed among the clusters identified based on the dbnSim matrix from RNAforester. While some RNA types form standalone clusters on their own, several others are split across multiple clusters, highlighting non-uniformities within individual types. In particular, the two least populous RNA types (TERC and 23S rRNA) are identified as outliers and clustered with other types. It is possible to have TERC and 23S rRNA as independent clusters with other clustering parameters allowing clusters with low densities, as shown in Fig. 5, though this leads to merging of tRNA and 5S rRNA and creation of many small clusters of the same RNA type. Taken all together, we can conclude that each RNA type represents a collection of most similar RNA molecules distant from all other types, while there exist substantial heterogeneities, both intra-type and inter-type, preventing a single set of algorithmic parameters from achieving perfect congruence between similarity-based clusters and RNA types. RNA types are thus used as cluster labels for this study.

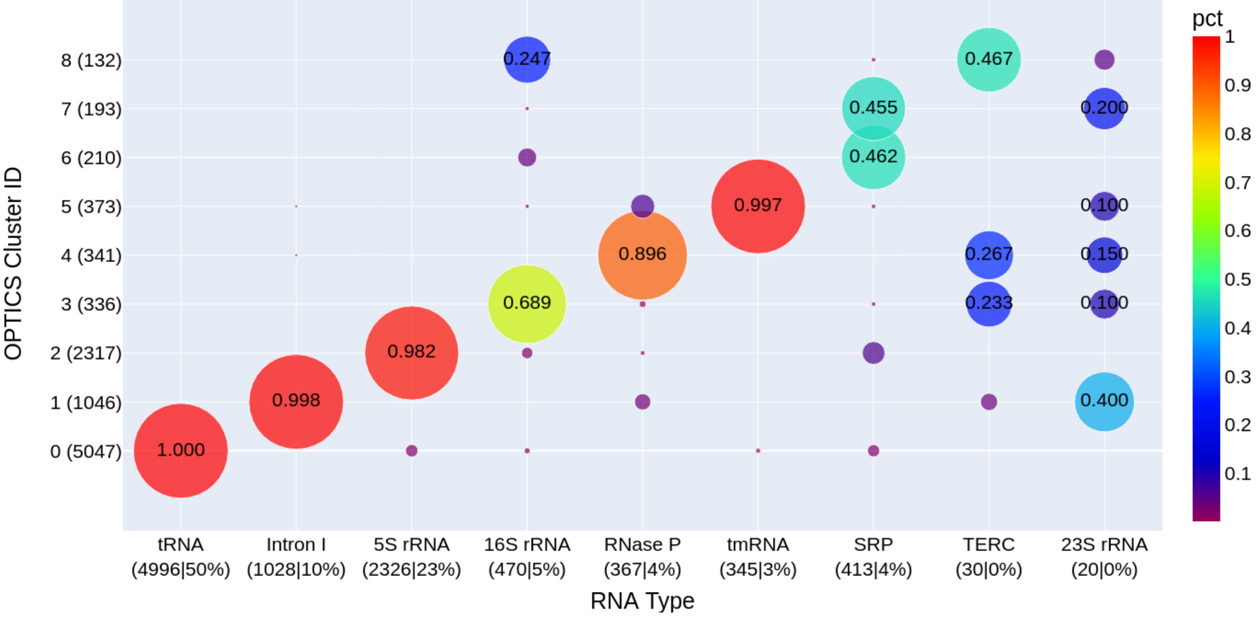


Figure 4 The distribution among similarity-based clusters for each RNA type. The size of each circle scales with the count of samples normalized by the total population of its respective RNA type, the numeric value of which is shown at the center of the circle.

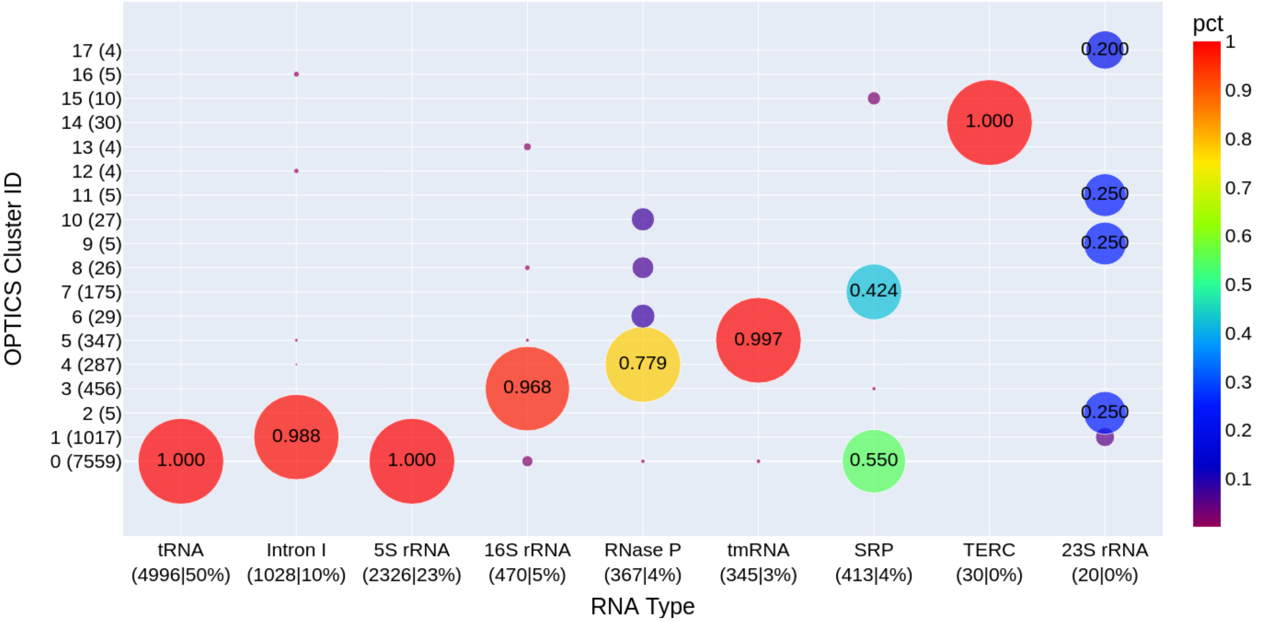


Figure 5 The distribution among similarity-based clusters for each RNA type as shown in Fig. 4. The difference is that the OPTICS parameters here allow clusters with lower densities and fewer samples, resulting in numerous clusters of rather small sizes.

### 2. SeqFold2D Model Architecture and Training

**Architecture.** The neural network (NN) architecture follows our previous work (Qiu, 2023). As shown in Fig. 6, it consists of two main modules operating on the base and pair representations in a serial manner. For an RNA sequence with length L, it is first embedded into an L×C matrix, where C is the number of channels, through one-hot encoding followed by a feed forward layer. The L×C matrix passes through a stack of N bi-directional Long-Short-Term-Memory (LSTM) blocks to update each base with its sequential context. Then, the pair representation matrix (L×L×C) is obtained via an outer-product transformation. The same N stack of 2D convolutional blocks is then employed to learn the local contact maps before the final feedforward layer to yield the logits for the base pairing probability matrix (PPM). Dropout and non-linear activation (ReLU) are applied after all linear transformations, except at the input and output. The total number of trainable parameters is determined by the choice of C and N. For this study, C is 64 and N is 4, resulting in ~960K parameters. All SeqFold2D models were implemented with the Paddle framework (https://github.com/PaddlePaddle/Paddle).

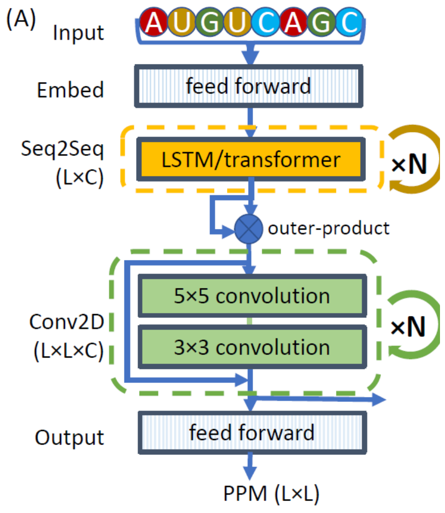


Figure 6 Network architecture of the SeqFold2D model, as shown in Fig. 1 in Ref. (Qiu, 2023)

**Training protocol.** Two loss functions are used: 1) Binary cross entropy (BCE) between the predicted PPM and the ground truth contact map (0 and 1 values only), and 2) Soft F1 score between the same pair of matrices. In binary classification, F1 score is defined as 2×Precision×Recall/(Precision+Recall)=2×TP/(2×TP+FP+FN), where Precision is TP/(TP+FP), Recall TP/(TP+FN), TP the number of true positives, FP false positives, FN false negatives, and TN true negatives. Here the soft F1 score is computed by weighing the 0 and 1 values in the contact map by the continuous probabilities in the PPM, instead of discrete counting. A typical training run starts with the BCE loss for about 50 epochs and then switches to the soft F1 score for about 20 epochs, with a learning rate of 0.001. For both loss functions, equal weights are applied to the 0 and 1 values in the ground truth. The AdamW optimizer is used for both loss functions.

### 3. Performance of LOCO SeqFold2D models

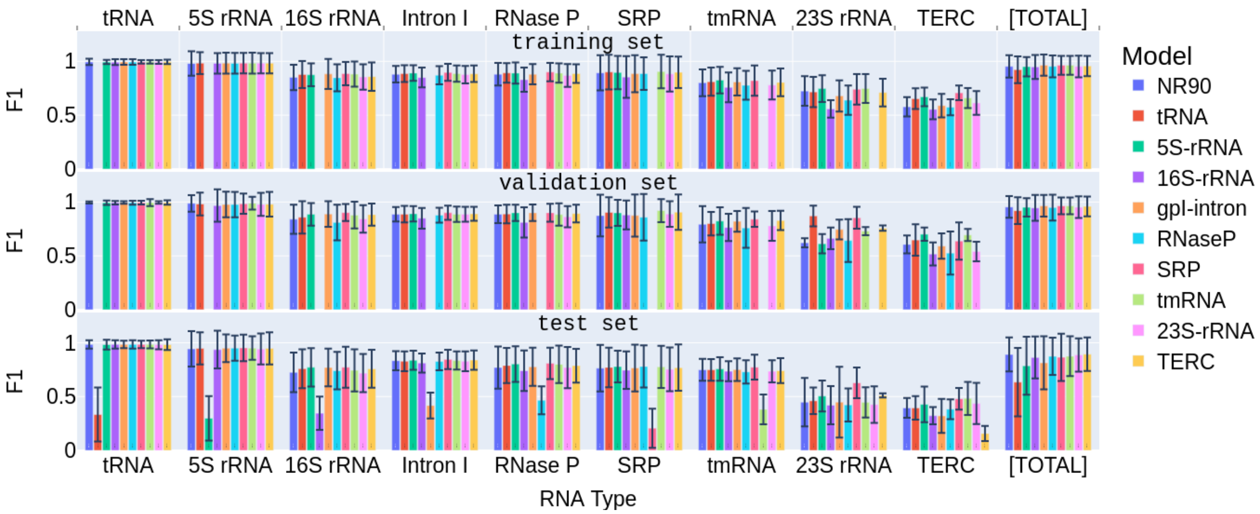


Figure 7 The performances of SeqFold2D models trained on the NR90 seen set. See the caption of Fig. 2 in the main text for detailed information.

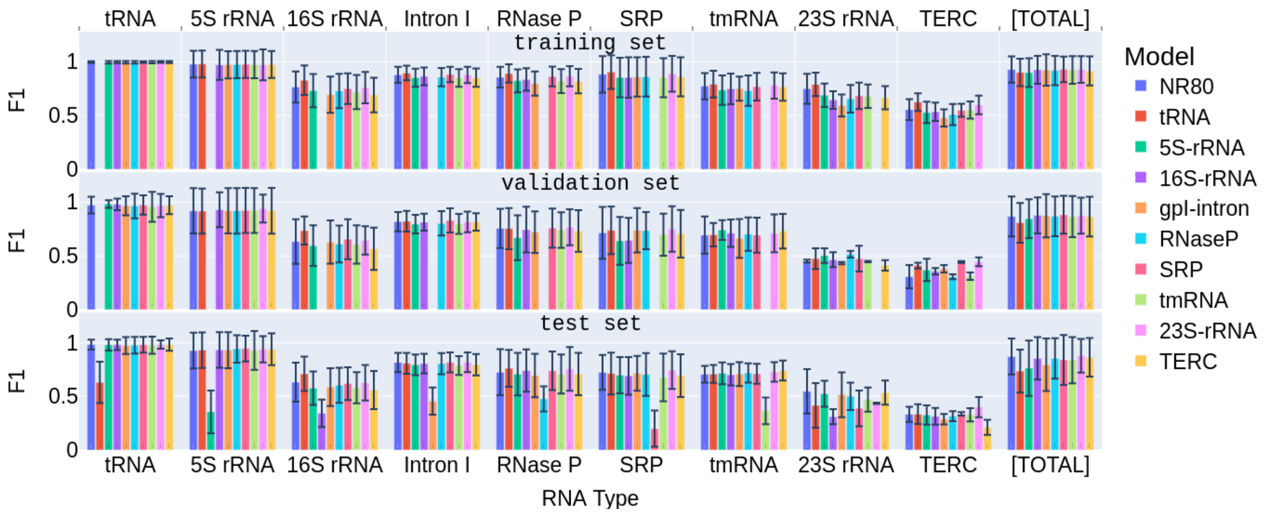


Figure 8 The performances of SeqFold2D models trained on the NR80 seen set. See the caption of Fig. 2 in the main text for detailed information.

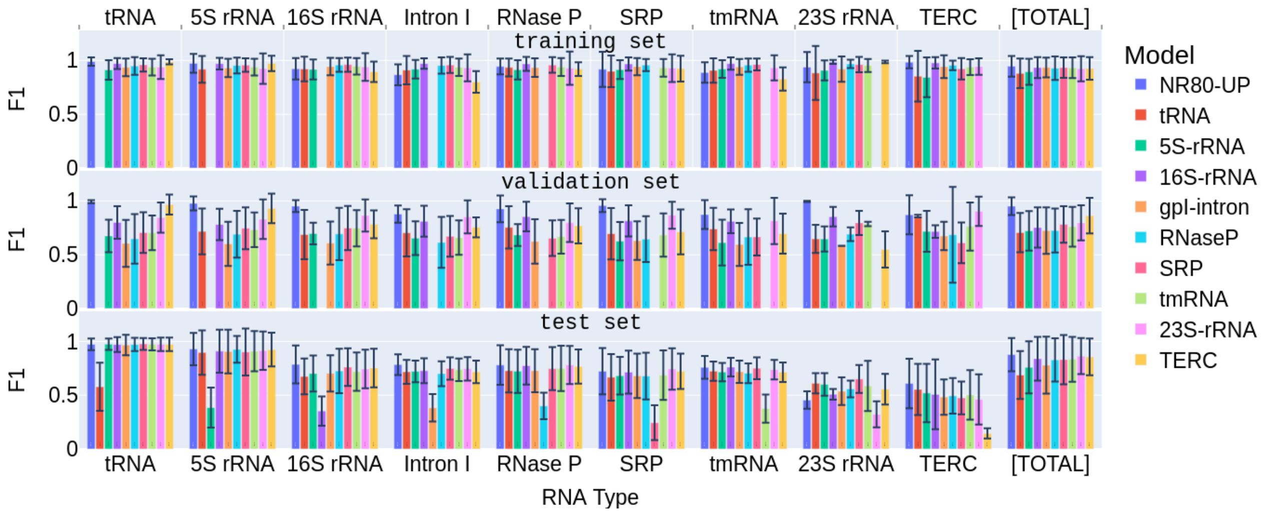


Figure 9 The performances of SeqFold2D models trained on the NR80-UP seen set. See the caption of Fig. 2 in the main text for detailed information.
